# Random UV Mutagenesis for the production of *Chlorella vulgaris* mutants with low chlorophyll content for the food industry

**DOI:** 10.1101/2025.04.28.651037

**Authors:** Ivan N. Ivanov, Jiří Kopecký, Karolína Štěrbová, Pavel Hrouzek, Martin Lukeš, Kateřina Bišová, Setyo Budi Kurniawan

**Affiliations:** Laboratory of Cell Cycles of Algae, Centre Algatech, Institute of Microbiology of the Czech Academy of Sciences, Novohradská 237, Třeboň, 379 81, Czech Republic; Laboratory of Algal Biotechnology, Centre Algatech, Institute of Microbiology of the Czech Academy of Sciences, Novohradská 237, Třeboň, 379 81, Czech Republic; Laboratory of Photosynthesis, Centre Algatech, Institute of Microbiology of the Czech Academy of Sciences, Novohradská 237, Třeboň, 379 81, Czech Republic; Department of Chemical and Process Engineering, Faculty of Engineering and Built Environment, Universiti Kebangsaan Malaysia, UKM Bangi, Selangor 43600, Malaysia; Research Center for Environment and Clean Technology, National Research and Innovation Agency (BRIN), Jakarta Pusat 10340, Indonesia

**Keywords:** alternative food, lutein, microalgae, mutants, yellow algae

## Abstract

Microalgae is currently gaining attention as an alternative source for food production. The market is currently demanding colorless algae with high protein and lutein content as an alternative to currently available commodities. This research was aimed at performing random UV mutagenesis on *Chlorella vulgaris* to obtain mutants with enhanced growth rates and increased growth characteristics. A total of seven mutants were selected to be analyzed after the random mutagenesis. Small (40 mL) and larger-scale (1,000 mL) reactors were used to analyze the production of *C. vulgaris* mutant biomass, focusing on the dry matter, starch, chlorophyll, protein, fatty acids, and lutein contents. Results implied that mutants showed a higher specific growth rate (2.1–2.5-fold higher) as compared to the wild type. The three mutants (MT 1, 2, and 3) that exhibited a yellow color were subsequently chosen for further scalability. In larger-scale reactors, all mutants exhibited higher protein contents while displaying lower carbohydrate and chlorophyll contents in comparison to the wild type. Moreover, MT 1 exhibited the highest concentration of lutein (0.37%–0.38%) and the lowest concentration of chlorophyll (0.1–0.14%), both of which are of significance for potential applications in the food industry.

## 1. Introduction

The demand from consumers for nutritionally dense and health-promoting consumables has increased in recent years. Microalgae possess a balanced biochemical profile and are a sustainable biological resource; they are abundant in protein and bioactive compounds, including carotenoids and essential fatty acids, which have the potential to positively impact human health (Schüler et al., 2020). However, it is worth noting that only a limited selection of the thousands of microalgal strains that have been identified and described are presently deemed safe for human consumption. Established on the market and with a long history of safe use, *Arthrospira platensis* and *Chlorella vulgaris* are permitted for human consumption in the European Union (Nunes et al., 2023).

Microalgal biomass is extensively utilized in the nutraceutical industry as dietary supplements (Galasso et al., 2019). In the food industry, it is typically integrated into conventional food products (e.g., bars, pasta, and cookies) as a healthy supplement or as a natural food colorant (Lafarga, 2019). However, the integration of microalgae into food items encounters obstacles primarily attributable to their organoleptic attributes, such as a potent hue, flavor, and odor (Wu et al., 2023). Consumer acceptance is significantly influenced by the sensory attributes of foods. Among these attributes, color is the initial parameter that consumers perceive, and it can potentially determine their dietary inclusion of the food in question; thus, culinary products derived from microalgae, which are typically green in color, have a negligible sensory appeal to consumers (Matos et al., 2022). It is imperative to modify these unfavorable organoleptic attributes of microalgal biomass to enhance its acceptability in food products.

In an effort to enhance the sensory attributes of food products containing microalgal biomass, some alternative approaches that have been explored include the simultaneous extraction of the desired compounds and chlorophyll removal, as well as the incorporation of components like chocolate to augment the ultimate flavor and color (Galanakis, 2021; Lucas et al., 2018). Isolation of novel microalgal species exhibiting enhanced organoleptic properties could also be considered (Schüler et al., 2020). Random mutagenesis is a noteworthy instrument for manipulating cells in the context of food applications. Unlike other methods that introduce external genetic material into the target cell, it does not produce genetically modified organisms (Bleisch et al., 2022; Kanakdande et al., 2021; Trovão et al., 2022).

Mutation breeding is an important method for producing microalgae strains with improved biomass composition and high growth rates (Thurakit et al., 2022). Chemical mutagenesis (Thurakit et al., 2022) and ultraviolet (UV) irradiation in particular are two of the most commonly used mutation breeding techniques for inducing random mutations in the microalgae genome (Bleisch et al., 2022). One of the major benefits of UV mutagenesis, in comparison to chemical methods, is that UV mutagenesis is random (Trovão et al., 2022).

Moreover, UV mutagenesis can lead to achieving higher benefits due to the significant biological effects of the mutations caused by UV on the organism (Kanakdande et al., 2021). Another major benefit of UV mutagenesis in comparison with the chemical methods is the ease of application and the relative safety of the procedure (Bose, 2014). UV mutagenesis of microalgae has been successfully applied for the development of a variety of biotechnology relevant strains in a wide number of microalgal species including *Chlorella* sp. (Liu et al., 2015). *Chlorella vulgaris* in particular, has gained increasingly popularity in mutagenesis for the production of biomass, lipid, and fatty acids (Kim et al., 2020; Sarayloo et al., 2018; Sivaramakrishnan and Incharoensakdi, 2023; Smalley et al., 2020).

Although the development of some *C. vulgaris* mutants has already been conducted for biomass, lipid, and fatty acids productions, research on the mutants with low chlorophyll content and improved growth characteristics is currently limited. This research is aimed at performing random UV mutagenesis for the production of *C. vulgaris* mutants with low chlorophyl content and enhanced growth characteristics for further applications in the food industry. The presented results are expected to enrich the knowledge of the alternative food sourced from microalgae.

## 2. Materials and Methods

### 2.1 Microorganism and cultivating conditions

#### Chlorella vulgaris

strain R117 (CCALA 1107) was obtained from the culture collection of Autotrophic Organisms, Institute of Botany, Třeboň, Czech Republic (further abbreviated as WT). For the growth screening, an RTS-8 multi-channel bioreactor (Biosan, Latvia) with the following settings: culture volume of 40 mL, temperature of 30 °C, and 2,000 rpm agitation were used. For growth and biomass composition comparison a set of BioBench twin fermenter units (Biostream International BV, Netherlands) with working volume of 1 L was used. The cultivation parameters of the fermenters were set as follows: temperature – 30 °C, aeration with atmospheric air – 1000 L min^−1^ and stirrer speed – 300 rpm. The cultivation systems are presented in Figure 1. Unless specified otherwise, all experiments were performed using ½ ŠS medium (Hlavová et al., 2016) supplemented with 20 g L^−1^ of glucose under dark condition. All experiments were performed in biological triplicates.

**Figure 1.**
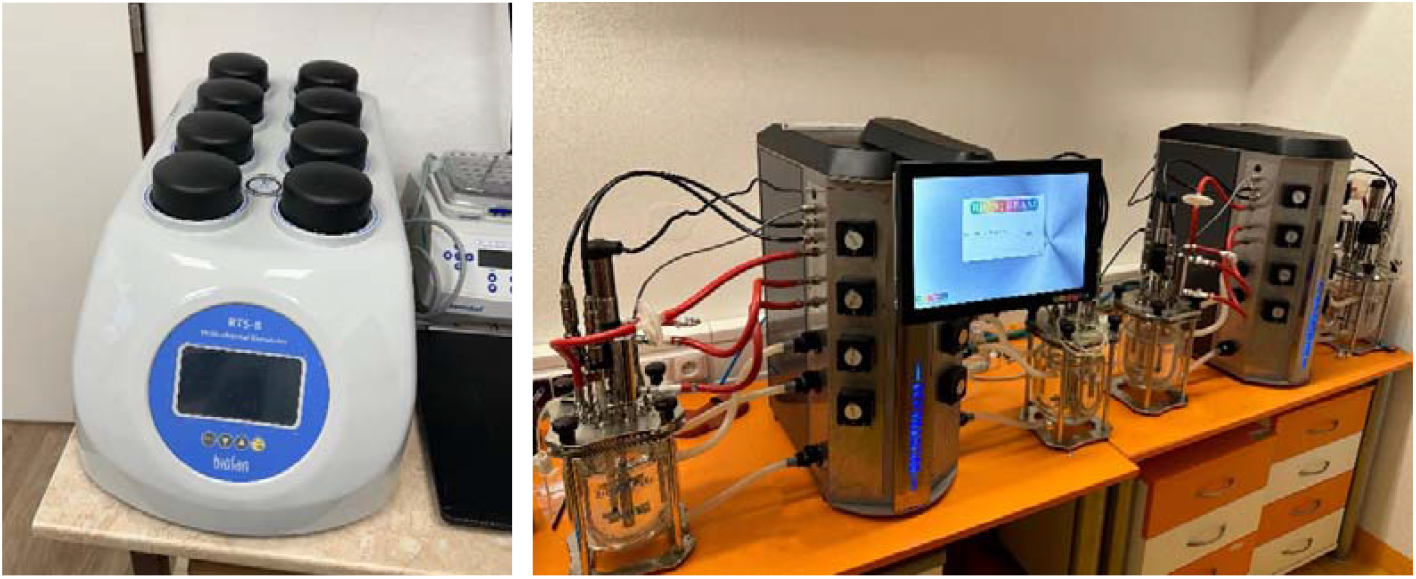
Cultivation systems: Biosan (left) and Biostream fermenters (right)

### 2.2 Mutagenesis and selection procedure

Cultures of *C. vulgaris* were grown in tubular bioreactors containing 300 ml ½ ŠS medium supplemented with 10 g L^−1^ glucose. The cultures were cultivated at a temperature of 30 °C and light irradiation of 500 µmol m^−2^ s^−1^ photosynthetically active radiation until an optical density at 750 nm (OD_750_) of approximately 1 was reached. After dilution to an OD_750_ of approximately 0.1, 25 mL of the cultures were poured into sterile glass Petri dishes with stirrers. The Petri dishes with the cultures were then irradiated with UV light for 5 minutes in a sterile hood. The irradiated cells were immediately transferred to sterile 100 ml flasks and incubated in the dark for at least 12 hours to avoid photoreparation. The cell number of the cultures was then counted and approximately 1,000 cells were seeded into a Petri dish containing ½ ŠS medium with 10 g L^−1^ glucose. The Petri dishes were incubated in the dark at 30 °C until colonies were visible. Cell survival was assessed by counting the number of colonies in the treated and control cultures. Only the non-green or pale green colonies were selected for further analysis.

### 2.3 Cell number measurements and growth assessment

For cell number analysis, 1 ml of the culture was fixed with 100 μL of 2.5% glutaraldehyde and stored at 4 °C until the time of analysis (Kselíková et al., 2022). Cell numbers were then analyzed using a Beckman Coulter Multisizer 4 (Beckman Coulter Life Sciences, Brea, CA, United States) by diluting 50 µL of the fixed cell suspension in 10 ml of 0.9% NaCl (w/v) electrolyte solution.

### 2.4 Dry matter measurements and growth assessment

At each sampling point, an aliquot of 2 mL culture suspension was taken and transferred to a pre-weighted (Sartorius TE214S-0CE, Germany) 2 mL plastic micro test tube that was pre-dried for 24 hours at 105 °C. The samples were centrifuged (Mikro 200, Hettich, USA) for 2 minutes at 14,000 g. The supernatant was removed and the test tube with the resulting pellet of cells was dried for 24 hours at 105 °C. After cooling down for at least 2 hours in a desiccator the test tube was weighted. The weight of the pellet was determined by subtracting the weight of the previously measured empty test tube (Kselíková et al., 2021).

### 2.5 Starch analysis

At each sampling point, 2 mL of algal suspension was centrifuged at 15,000 g and stored at −20 °C until use. The quantity of starch was determined using the approach by McCready et al. (1950), with the exception of the extraction procedure, which is detailed in the following section. The algal cells were disintegrated by vortexing with 300 µL of glass beads in 500 µL of distilled water. Pigments were extracted three times in 1 mL of 80% ethanol at 68 °C for 15 minutes. Total starch within the whitish pellet was hydrolyzed by 1.5 mL of 30% perchloric acid followed by agitation for 15 min at room temperature. Following centrifugation of the samples, the supernatant was extracted. The procedure was repeated three times to yield 4.5 mL of hydrolyzed starch extract; this volume was subsequently diluted to a final volume of 5 mL (Kselíková et al., 2022).

### 2.6 Total chlorophyll analysis

An aliquot of 10 mL culture suspension was taken and transferred to a 15 mL falcon tube which was centrifuged for 10 min at 2772 g (Rotina 380R, Hettich, USA) and stored at −20 °C. For the disintegration of the cells, 1 mL of phosphate buffer, pH 7.7, and 500 μL of glass beads were added to the pellet. The samples were vortexed vigorously for 10 minutes at room temperature(Vortex Genie 2, Scientific Industries, Inc., NY, USA). The samples were extracted with 4 mL of 100% acetone, briefly vortexed and centrifuged for 3 min at 4332 g. The supernatant was transferred into graduated glass test tubes with a stopper and put into a dark block. The pellet was again extracted by 4 mL of 80% acetone followed by brief vortexing and centrifugation (3 minutes, 4332 g); the supernatant was transferred into the same glass tube as before. The two combined extracts in the glass tubes was filled up to 10 mL with 80% acetone. The absorbance of the samples was measured in a spectrophotometer (UV-1900i, Shimadzu, Japan) at the following wavelengths: 750, 664, 647, 470 and 450 nm. The total chlorophyll concentration was calculated according to the equations and coefficients for 80% acetone in Wellburn, 1994 (Wellburn, 1994).

### 2.7 Fatty acids analysis

The fatty acids analysis was performed according to Ranglová et al. (2022). The samples were collected throughout the cultivation and an amount corresponding of approximately 5 to 10 mg of freeze-dried biomass was used for the analysis.

### 2.8 DNA and protein analysis

#### 2.8.1 Total Nucleic Acids

The procedure of Wanka (1962) as modified by Lukavský et al. (1973) was used for the extraction of the total nucleic acids. The samples were centrifuged in 10 mL centrifuge tubes, which also served for storage of the samples. The pellet of algal cells was stored under one ml of ethanol at −20 °C. The pellet was extracted 5 times with 0.2 M perchloric acid in 50% ethanol for 50 min at 20 °C and 3 times with an ethanol-ether mixture (3:1) at 70 °C for 10 min. Such pre-extracted samples can be stored in ethanol at −20 °C. Total nucleic acids were extracted and hydrolyzed by 0.5 M perchloric acid at 60 °C for 5 h. After hydrolysis, concentrated perchloric acid was added to achieve a final concentration of 1 M perchloric acid in the sample. Absorbance of total nucleic acids in the supernatant was read at 260 nm (A260) and total nucleic acid concentration was calculated based on the calibration with DNA standard of the known concentration treated by the same procedure.

#### 2.8.2 DNA and RNA Determination

The light activated reaction of diphenylamine with hydrolyzed DNA, as described by Decallonne and Weyns (1976) was used with the following modification of Zachleder (1984): the diphenylamine reagent (4% diphenylamine in glacial acetic acid, w/v) was mixed with the samples of the total nucleic acid extracts in a ratio of 1:1 and the mixture in the test tubes was illuminated from two sides with fluorescent lamps (L 40W F33, Tungsram, Budapest, Hungary). The incident radiation from each side was 150 µmol photons m^−2^ s ^−1^. After 6 h of illumination at 40 °C, the difference between the A600 and A700 nm was estimated. The concentrations of DNA within the samples were set by comparison to the A600 and A700 nm values of the sample with known DNA concentration treated the same way. The values were normalized to the number of cells extracted. The RNA content was calculated as the difference between the total nucleic acid and DNA content.

#### 2.8.3 Protein Determination

The pellet remaining after nucleic acid extraction was used for protein determination. It was hydrolyzed by 1 M NaOH for 1 h at 70 °C. The protein concentration in the supernatant after centrifugation of the hydrolysate (15 min, 5300× g, room temperature) was estimated by BCA assay (cat. no. 23225, Thermo Fisher Scientific, Waltham, MA, USA) according to manufacturer’s conditions. The same procedure was carried out with a calibration curve set by different concentrations of bovine serum albumin.

### 2.9 Lutein extraction and HPLC analysis

A 2 mL aliquots of the culture were sampled and transferred to 2 mL plastic micro test tubes. The plastic test tubes were then centrifuged at 13,148 g for 2 min. (Minispin plus, Eppendorf SE, Germany) after which the supernatant was removed. For storage purposes the test tubes with the resulting pellets of microalgae biomass were placed at −20 °C. A total of 2 mL of the extraction solvent (55.4% (v/v) H_2_O, 44.5% (v/v) ethanol, 0.1% (v/v) n-heptane) was added to each pellet after which the samples were put in an ultrasonic bath for 5 min (Kranitek 6, Kranitek Czech s.r.o. Czech Republic). After that, the samples were centrifuged for 5 min at 1,073 g. The supernatant was transferred to a graduated glass test tube after which the extraction step was repeated 1 or 2 more times, depending on the amount of algal biomass.

The total volume of solvent used for each sample was noted and was used later on for the calculation of the lutein concentration. A total of 1 mL of supernatant was taken from each graduated glass test tube and was transferred into a glass HPLC vial. The lutein concentration of the samples was measured on an HPLC (Agilent 1100 Series, Agilent Technologies GmbH, Germany) equipped with aLuna C8(2) HPLC column (Phenomenex, Torrance, CA, USA).

### 2.10 Statistical analysis

All experiments were performed in at least three biological replicates. If not stated otherwise, all results are presented as means and standard deviation (n = 3).

## 3. Results and discussions

### 3.1 Mutagenesis

The cultures of *Chlorella vulgaris* were grown mixotrophically to promote expression of genes required for both organic carbon utilization and for photosynthesis. Exponentially growing culture were diluted 10-fold and used directly for mutagenesis by UV light. Different durations of UV light irradiation were tested to produce mutant populations. As expected, the survival rates were decreasing with prolonged irradiation. While at irradiation for 1 and 2 minutes the survival rates ranged between 74-93 %, they dropped to approximately 30 % after 4 minutes and to 12 % after 5 minutes of irradiation. Survival rate between 10-15 % was selected as a rate producing enough mutations but not affecting the cell cultures detrimentally. Approximately 60,000 dark grown mutants were colour selected yielding 7 mutants (MT 1 to 7) with lighter colors ranging from yellow to light green.

### 3.2 Biosan experiments: growth and biomass analysis

The seven mutants were characterized in detail in small volume (40 ml) cultures grown heterotrophically in mineral medium (1/2 ŠS) containing 2% glucose at 30 °C. The wild type (WT) reached maximum cell concentration of 52 × 10^6^ cells mL^−1^; the mutants varied between 44 × 10^6^ cells mL^−1^ and 689 × 10^6^ cells mL^−1^ (Table 1). Two of the mutants, MT 1 and MT 2 reached significantly higher maximum cell concentrations than wild type, 628 and 689 × 10^6^ m L^−1^, i.e. 12- and 13-fold higher than the wild type. However, the maximum OD reached was similar between wild type and MT 1 and 2. This together with the cell sizes and dry matter per cell values suggests the only difference might be in the speed of the final cell division (Table 1). Specific growth rates (µ) of all the mutants (ranging from 0.28 to 0.33 h^−1^) were higher than that of wild type (0.13 h^−1^), consequently the doubling times of the mutants were shorter (ranging from 2.1 h to 2.45 h) than that of wild type (5.33 h). The specific growth rate of mutants reached 2.1 to 2.5 fold higher as compared to WT, which is higher as compared to previous research by Ong et al. (2010) which reported 1.4 – 1.8 fold higher growth rate of *Chlorella* sp. and Kim et al. (2020) which mention 1.3 fold higher growth of *C. vulgaris* mutant as compared to the wild type.

**Table 1.** Growth characteristics of the mutant strains.

| Strain | Maximum cell concentration (10<sup>6</sup> cells ml<sup>-1</sup>) | Maximum OD | Specific growth rate $\mu$ (h<sup>-1</sup>) | Doubling time (h) |
| --- | --- | --- | --- | --- |
| WT | 52.06 ± 2.47 | 40.66 ± 0.82 | 0.13 ± 0.07 | 5.33 |
| MT 1 | 627.7 ± 44.23 | 51.1 ± 2.73 | 0.29 ± 0.38 | 2.39 |
| MT 2 | 688.9 ± 75.18 | 56.16 ± 2.15 | 0.28 ± 0.37 | 2.45 |
| MT 3 | 70.3 ± 2.94 | 13.26 ± 0.23 | 0.29 ± 0.38 | 2.39 |
| MT 4 | 101.3 ± 27.8 | 21.99 ± 0.07 | 0.33 ± 0.37 | 2.1 |
| MT 5 | 55.4 ± 0 | 54.05 ± 0 | 0.29 ± 0 | 2.39 |
| MT 6 | 53 ± 6.16 | 30.44 ± 26.43 | 0.29 ± 0.34 | 2.42 |
| MT 7 | 44.16 ± 2.51 | 10.32 ± 0.37 | 0.28 ± 0.37 | 2.45 |

The cell composition analyses (Table 2) showed no difference in the proportion of DNA and RNA per dry matter (DM) or cell; there were some minor differences in the starch content.

**Table 2.**
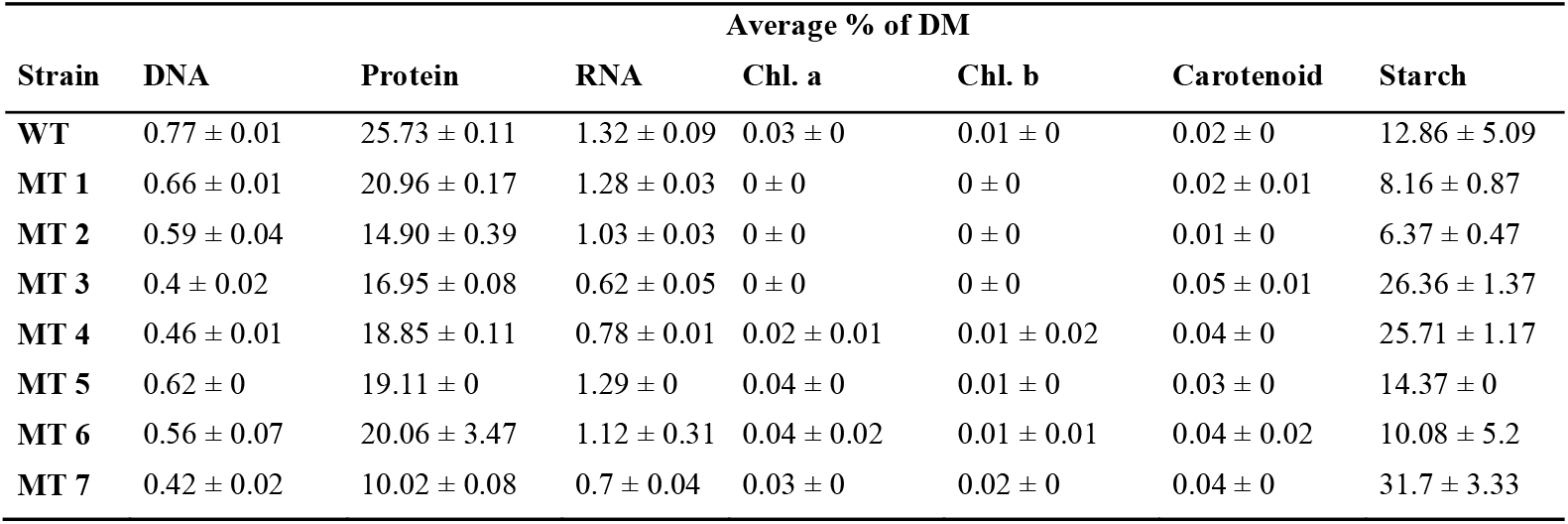
Biomass composition of mutant strains compared to the wild type.

The cell composition analysis confirmed low chlorophyll concentration in mutants no. 1-3 (depicted in Figure 2), where chlorophyll *a* and *b* were below detection limit. Low chlorophyll species of algae is currently gaining attention for further application in food industries (Schüler et al., 2020), which make these 3 mutants interested to be scaled up further. In mutants no. 4-7, there was no significant decrease in chlorophyll content despite the visual change in color. Fatty acid profiles did not show any differences between the wild type and mutants in either saturated or unsaturated FA. Some mutants (MT 3, 4, 5 and 7) exhibited higher starch content as compared to WT, which can be related to the accumulation of some polysaccharides under heterotrophic conditions (Schüler et al., 2020).

**Figure 2.**
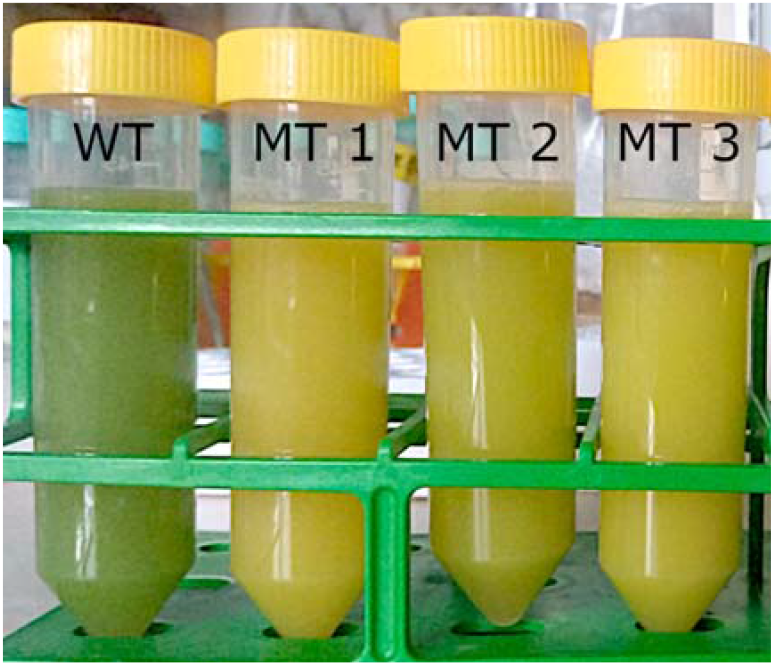
Differences in colouration of cultures of the wild type and mutant strains 1, 2, 3 grown in the RTS-8 multi-channel bioreactor after 50 hours.

### 3.3 Biostream cultivations: growth and biomass analysis

Given the low chlorophyll concentration and high growth rates, the mutant strains no. 1-3 were tested in larger scale, which confirmed the results from a small scale testing. Of all the strains, no. 1 and 2 seem to be most promising, particularly due to improved growth rates. Dry matter, OD, and cell number obtained during the Biostream cultivation are depicted in Figure 3, while calculation of growth rate and doubling time based on OD (28-40 h for WT, 44-48 h for MT1, 36-44 h for MT2, 44-52 h for MT3), are tabulated in Table 3. Even the obtained growth rate was lower as compared to the Biosan system, mutants maintained a higher value as compared to the WT. The MT 1 and 3 reached 1.6 fold higher specific growth rate than WT, similar to some previously published reports (Kim et al., 2020; Ong et al., 2010). In addition, mutants also maintained lower doubling time and higher dry matter after cultivation period (maximum of 4.43 g L^−1^ with 3.3 fold higher than WT).

**Table 3.** Growth characteristics of the wild type and the mutant strains cultivated in the Biostream fermenters. Data are shown as average values and standard deviation (n=4).

| Strain | Max. DM<br>(g L<sup>-1</sup>) | Max. OD<sub>750nm</sub><br>(r.u.) | Specific growth rate<br>$\mu$ (h<sup>-1</sup>) | Doubling time<br>h |
| --- | --- | --- | --- | --- |
| WT | 1.36 ± 0.10 | 4.54 ± 0.32 | 0.07 ± 0.01 | 10.40 ± 1.02 |
| MT 1 | 3.62 ± 0.44 | 13.32 ± 1.37 | 0.11 ± 0.03 | 6.77 ± 1.60 |
| MT 2 | 4.31 ± 0.44 | 14.22 ± 0.37 | 0.07 ± 0.00 | 9.72 ± 0.57 |
| MT 3 | 4.43 ± 0.24 | 15.82 ± 0.83 | 0.11 ± 0.00 | 6.51 ± 0.17 |

**Figure 3.**
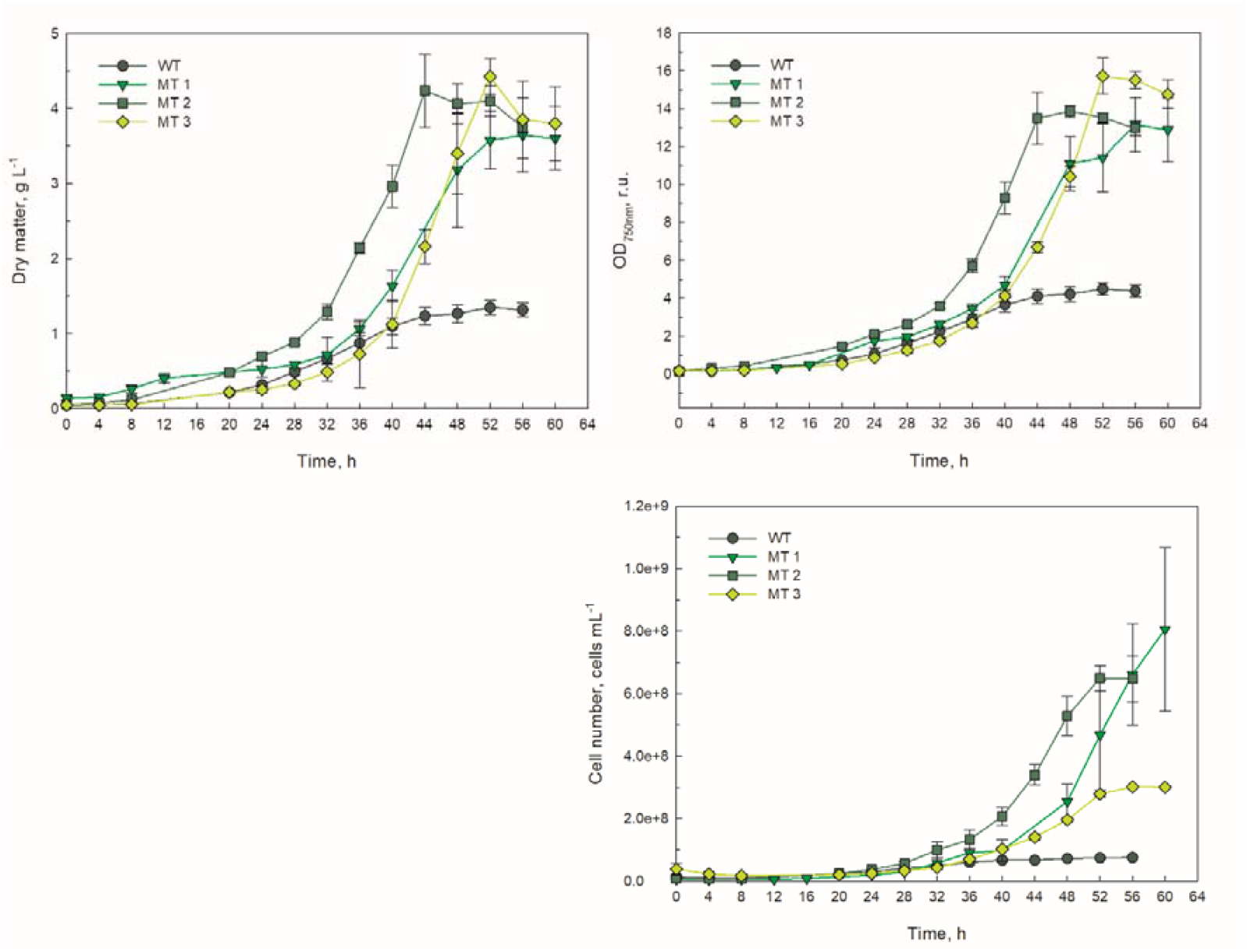
Dry matter, OD, and cell number of the wild type and the three mutant strains during cultivation in the Biostream fermenters. Data are shown as average values and standard deviation (n=4).

As presented in Figure 2, the yellow colour appeared on the MT 1, 2, and 3 cultures is subjected to the presence of xanthophyll lutein (Huang et al., 2018). The decreasing of chlorophyll content accompanied with the increase of lutein are interesting for food scientist since some microalgae-based food has low consumer sensorial acceptance due to the appearance of green colour (van Lelyveld and Smith, 1989). The chlorophyll content in WT was decreased along the test period while fluctuated for all mutants (Figure 4 and Table 4), with WT showed higher value than mutants as can be seen also by greenish colour in Figure 2. The decrease in chlorophyll content in WT indicate that the cell did not invest in chlorophyll biosynthesis under heterotrophic condition, while also indicating the degradative chlorophyll reactions (Beisel et al., 2010).

**Table 4.**
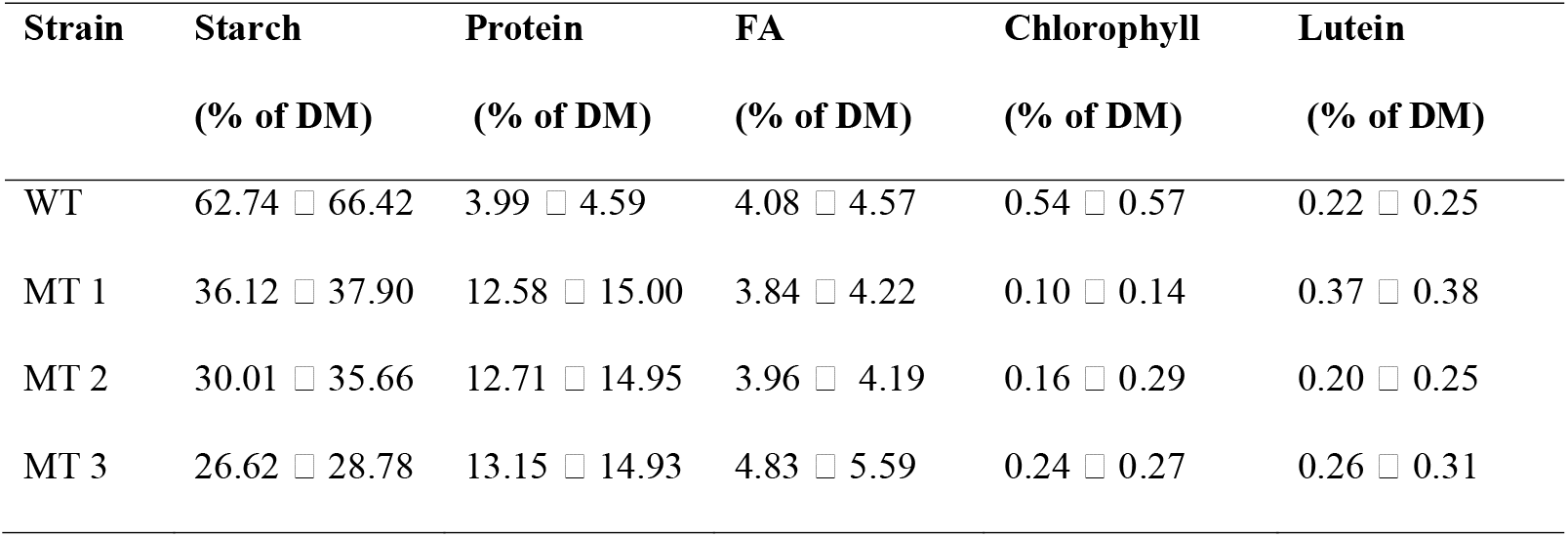
Range of macronutrients and pigments composition of the wild type and the mutants during the stationary phase.

**Figure 4.**
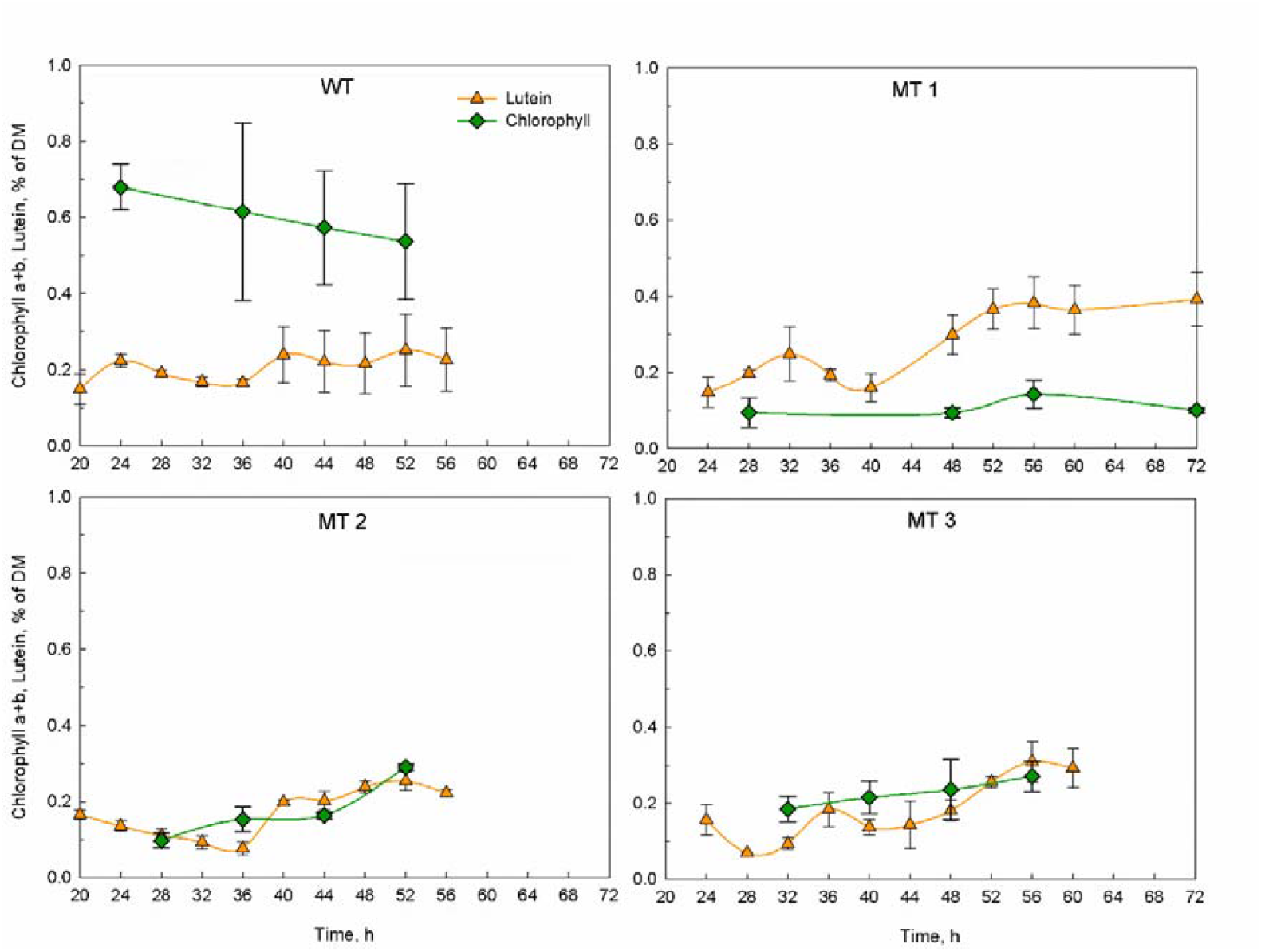
Chlorophyll a+b and lutein content of the wild type and the three mutant strains during cultivation in the Biostream fermenters. Data are shown as average values and standard deviation (n=4).

Referring to the starch content (Figure 5 and Table 4), WT showed a trend of increasing starch content which may be referred to the starch accumulation under heterotrophic conditions (Schüler et al., 2020). A decrease in protein and fatty acids content confirmed that starch accumulation was a mechanism of microalgae to prevent energy loss by reducing protein and lipid synthesis (Shi et al., 2022). Different from the wild type, decreasing trend was obtained for starch accumulation in all mutants, along with the increasing of protein (Figure 5 and Table 4). This suggests that mutants consumed internal carbohydrate (as the product of heterotrophic biosynthesis) to synthesize protein (Li et al., 2008), hence reducing the starch content in biomass. The observation of increased protein levels and decreased chlorophyll concentrations in mutant species may indicate that the photosystems have truncated chlorophyll antennas, called chlorophyll-binding proteins and thylakoid membrane proteins, as has been documented in other mutants lacking chlorophyll (Dall’Osto et al., 2019; Shin et al., 2016).

**Figure 5.**
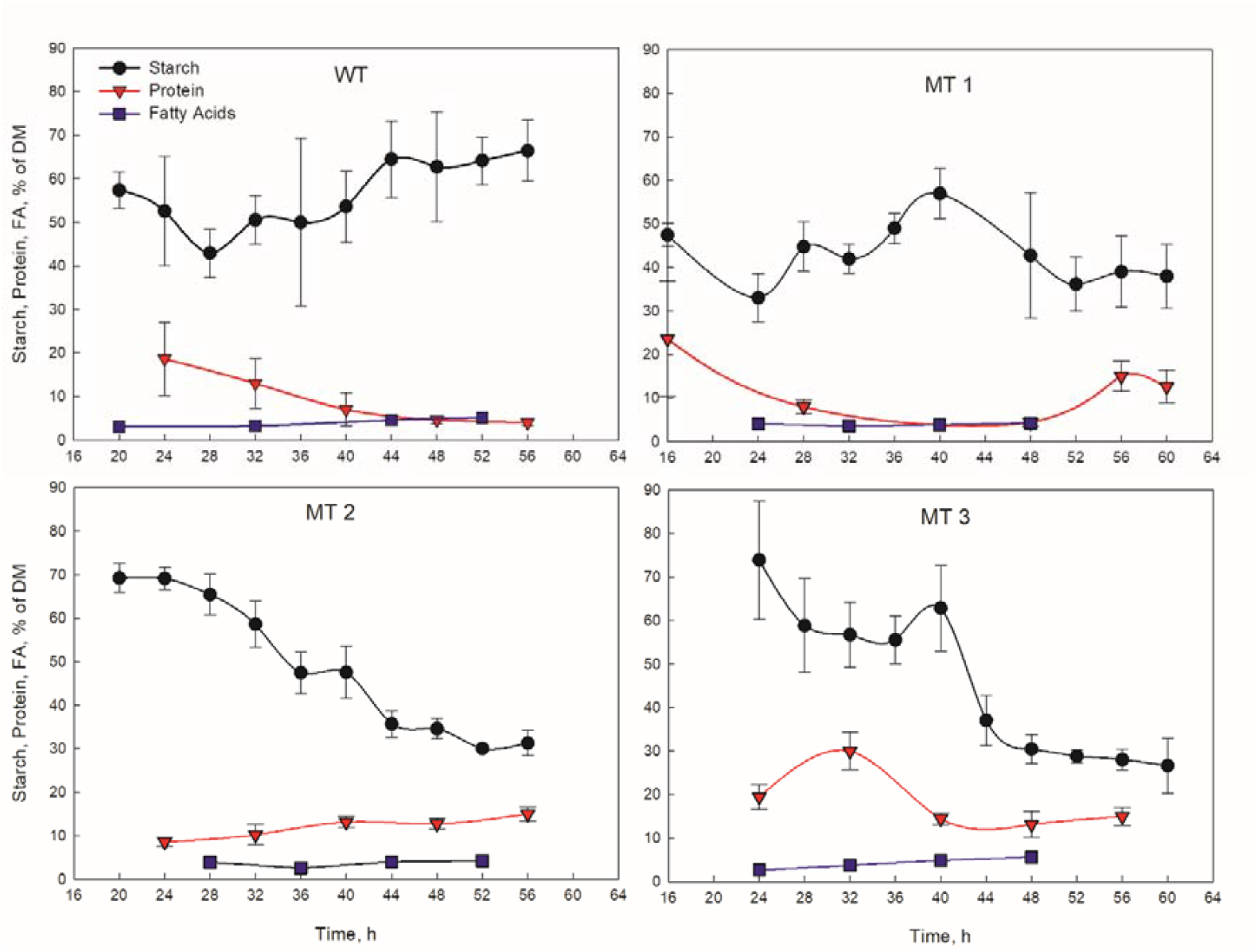
Biomass composition of the wild type and the three mutant strains during cultivation in the Biostream fermenters. Data are shown as average values and standard deviation (n=3 for starch and protein content and n=1 for fatty acids).

Despite the differences found in the starch accumulation, there is no significant difference found in the content of lipid and fatty acids between WT and mutants (Figure 5 and Table 4). The obtained result was in accordance with Schüler et al. (2020) which also reported the insignificant differences of lipid and FA content between WT and mutants. The fraction of TFA was also varied in each species (Figure 6), while C16:0 (palmitic acid) and C18:2 (linoleic acid) mostly composes the whole distribution (>20% of DW each). Both palmitic and linoleic acid are major constituents of TFA in the *C. vulgaris* as also stated previously by (Moradi-Kheibari et al., 2022).

**Figure 6.**
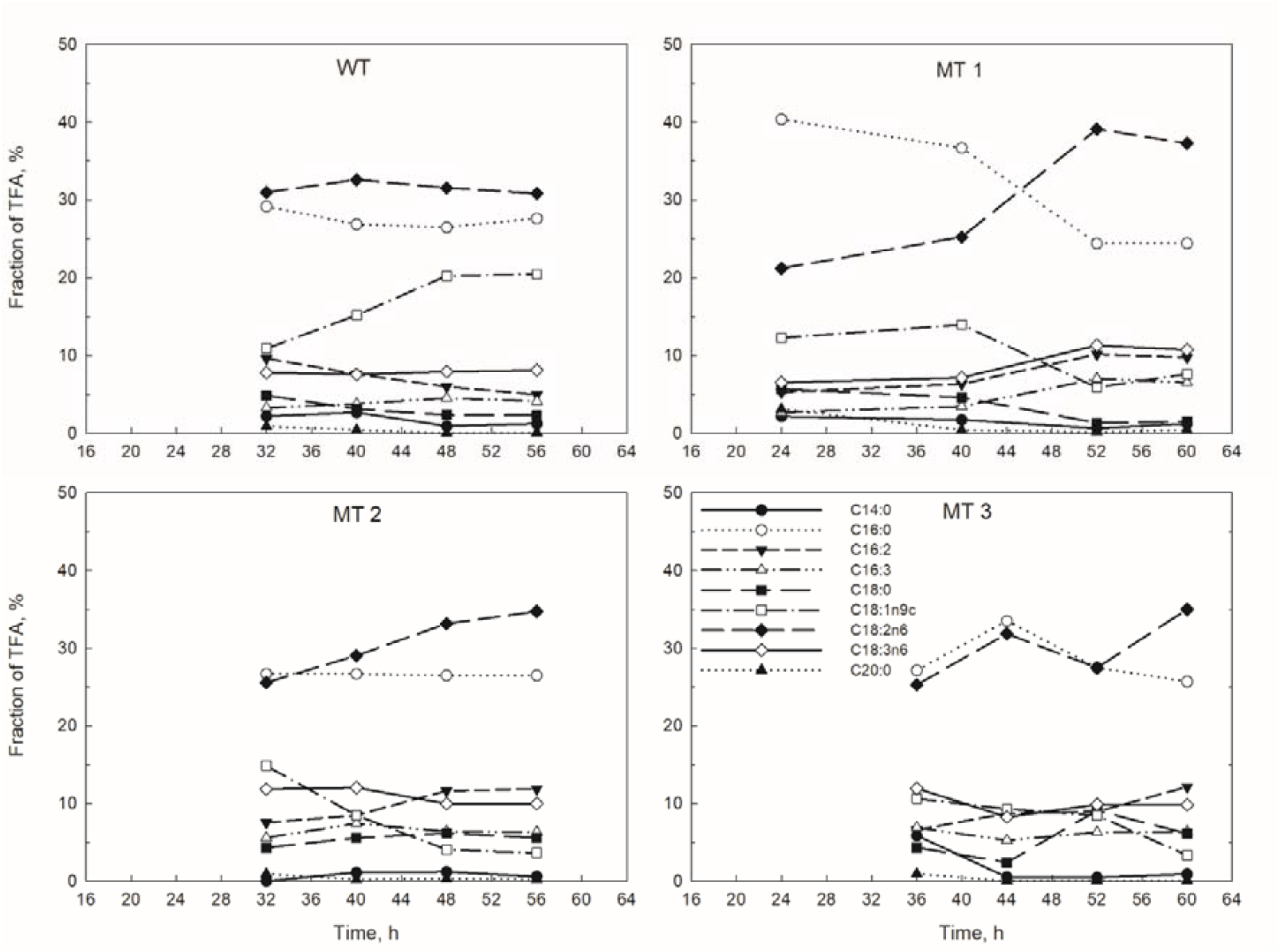
Fraction of total fatty acids in the wild type and the three mutants during cultivation in the Biostream fermenters (n=1)

## 4. Conclusion

Random UV mutagenesis to *Chlorella vulgaris* showed a good effect in term of chlorophyll, protein, and lutein contents. Based on the obtained results, all mutants showed higher specific growth rate, ranging from 2.1 to 2.5-fold, as compared to the wild type. Three mutants (coded as MT 1, 2, and 3) exhibited low chlorophyll concentrations, which then selected to be scaled up further. All mutants showed higher protein contents, lower starch and chlorophyll contents as compared to wild type in bigger scale reactors. Furthermore, MT 1 showed the highest lutein content 0.37 – 0.38% and lowest chlorophyll content 0.1 – 0.14%, which is interesting for further utilization in food industries.

## 5. Acknowledgements

This project has received funding from the Bio Based Industries Joint Undertaking (JU) under grant agreement No 887227. The JU receives support from the European Union’s Horizon 2020 research and innovation programme and the Bio Based Industries Consortium. For the purpose of Open Access, a CC-BY public copyright licence has been applied by the authors to the present document and will be applied to all subsequent versions up to the Author Accepted Manuscript arising from this submission.

